# Integrative Analysis Reveals Histone Demethylase LSD1/KDM1A Associates with RNA Polymerase II Pausing

**DOI:** 10.1101/2020.10.13.338103

**Authors:** Hani Jieun Kim, Taiyun Kim, Andrew J. Oldfield, Pengyi Yang

## Abstract

RNA polymerase II (RNAPII) pausing at gene promoters is a rate-limiting step in transcription regulation. Previous studies have elucidated the coordinated actions of pausing and releasing factors that collectively modulate RNAPII pausing. In general, the involvement of chromatin remodellers in RNAPII pausing has not been well documented. Whilst LSD1 is well-known for its role in decommissioning enhancers during ESC differentiation, its role at the promoters of genes remains poorly understood despite their widespread presence at these sites. Here, we report that LSD1 is associated with RNAPII pausing at the promoter-proximal region of genes in mouse embryonic stem cells (ESCs). We demonstrate that the knockdown of LSD1 preferentially affects genes with higher RNAPII pausing than those with lower pausing and, importantly, show that the co-localization of LSD1 and MYC, a factor known to regulate pause-release, is associated with the enrichment of other RNAPII pausing factors compared to their independent counterparts. Moreover, we found that genes co-occupied by LSD1 and MYC are significantly enriched for housekeeping genes that are involved in metabolic processes and globally depleted of transcription factors compared to those bound only by LSD1. These findings reveal a pleiotropic role of LSD1 in regulating housekeeping program besides its previously known role in regulating cell identity programs. Our integrative analysis presents evidence for a previously unanticipated role of LSD1 in RNAPII pausing through its association with pause release factors in modulating cell-type specific and cell-type invariant genes.

## Introduction

Pausing of the RNA polymerase II (RNAPII) during early elongation is an obligate part of the transcription cycle that is experienced by RNAPII at almost all genes^1^. Blocking the release of the paused RNAPII through the use of potent small molecule inhibitors has been shown to globally trap RNAPII at promoters, thereby abrogating nearly all RNA synthesis in mammalian cells^2–4^. Release of the paused RNAPII in promoter-proximal regions has been shown to be the rate-limiting step in productive elongation, making the efficiency of pause release a central determinant of gene expression. While early studies have shown that paused RNAPII is preferentially found at many developmental genes in the Drosophila melanogaster embryo^5^, subsequent research has demonstrated that pausing is widespread in higher eukaryotes and are not only limited to developmental genes but also genes associated with essential biological processes such as cell proliferation and stress response^6^. For example, in embryonic stem cells (ESCs), RNAPII pausing has been implicated in key processes such as self-renewal^7^ and differentiation^1, 2, 8, 9^. Thus, the identification of factors that modulate RNAPII pausing and release at genes of different functions is of great interest towards understanding gene regulation.

Many events and factors are involved in the establishment and release of paused RNAPII^10–13^. After recruitment to the promoter region of a gene, RNAPII begins transcribing a short, nascent RNA approximately 25-50 nucleotides long. However, RNAPII halts within the initially transcribed region and remains promoter-proximally paused until it receives further signals. The paused state is stabilized by SPT5 and NELF-A^14^, with NELF-A preventing reactivation of the RNAPII catalytic site^15–17^. The release of RNAPII into productive RNA synthesis is triggered by the activity of the kinase, positive transcription elongation factor b (P-TEFb), where CDK9 acts as the catalytic subunit of P-TEFb. Recruitment of P-TEFb was proposed to be facilitated by c-MYC (MYC), BRD4, various subunits of the Mediator and super elongation complexes^3, 18, 19^. Critically, the phosphorylation of SPT5 by P-TEFb has been shown to be directly linked to pause release^20–23^, causing the dissociation of NELF-A from RNAPII to enable reactivation and continued elongation of the nascent RNA^21, 24, 25^.

Here, we report the histone lysine-specific demethylase 1 (LSD1)^26^, a lysine-specific histone demethylase (also known as KDM1A/AOF2) which acts on mono- and dimethylated H3K4 and H3K9, as a putative factor associated with RNAPII pausing. LSD1 is the only demethylase to use a FAD-dependent oxidation reaction and specifically demethylates H3K4 and H3K9^26^. In ESCs, the inhibition of LSD1 has been shown to lead to severe proliferative defects and cell death when its activity is blocked during differentiation^27, 28^. Chromatin immunoprecipitation sequencing (ChIP-seq) analyses of LSD1 with other DNA/chromatin binding proteins in mouse revealed its essential role in decommissioning enhancers during ESC differentiation^27^. Whilst LSD1 was found to bind both gene promoters and enhancers in ESCs, the function of LSD1 has so far been focused on its action at enhancers^27, 29, 30^, and the function of its binding at gene promoters remains largely unknown.

We found that LSD1 presence is pervasive at gene promoters and in particular co-localizes with a number of factors associated with paused RNAPII. Amongst the RNAPII factors, it demonstrated the strongest co-localization with the release factor P-TEFb but also showed strong co-localization with another RNAPII pausing-releasing transcription factor, MYC. We show that the co-localization sites of LSD1 and MYC is associated with significantly higher RNAPII pausing, more enriched with pausing and releasing factors, and most sensitive to LSD1 knockdown. Furthermore, LSD1 targets different gene categories whether it binds independently of MYC or not. Collectively, our results implicate LSD1, a lysine-specific histone demethylase, in the regulation of RNAPII pausing and thereby gene expression and highlight a role of LSD1 at promoters that may be distinct from its role in ESC differentiation.

## Results

### LSD1 Co-localizes with Key Components of the RNAPII Pausing Machinery at Gene Promoters

We previously developed an analytic tool, PAD (http://pad2.maths.usyd.edu.au/), for identifying TFs, chromatin remodellers and histone modifications that co-localize on chromatin based on their genomic binding profiles, as measured using ChIP-seq^31^. Here, we extended PAD to enable targeted co-localization analysis within 12 functional genomic regions identified by ChromHMM^32^ (Fig. 1; see Methods).

**Figure 1.**
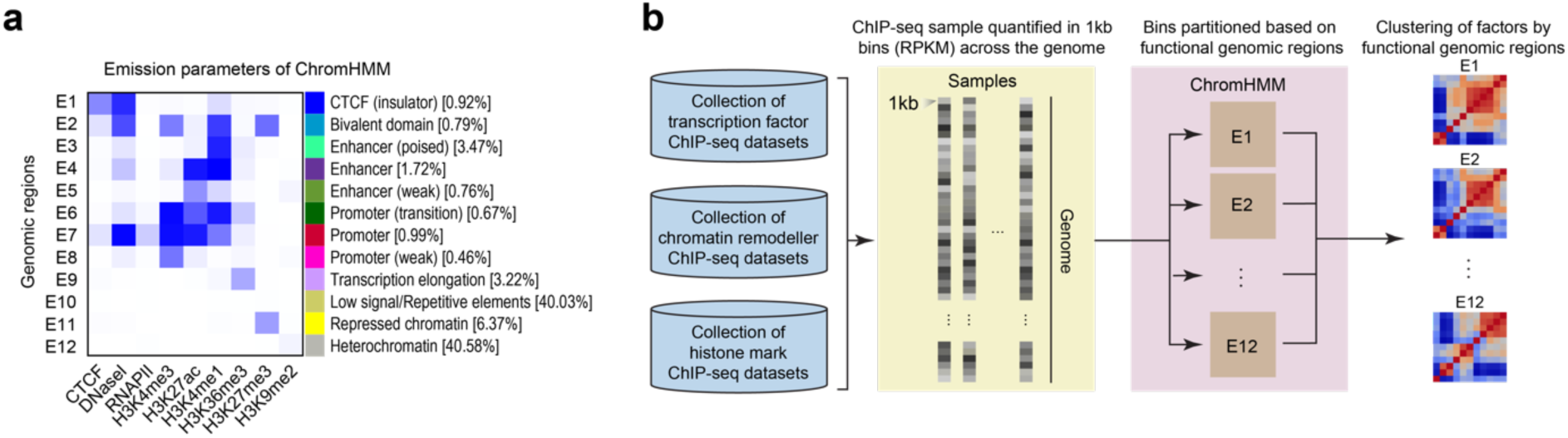
Implementation of PAD and evaluation of measures of ChIP-seq signal correlation. (a) A heatmap of functional genomic regions (y-axis) identified by enrichment of key histone markers and DNA and chromatin binding proteins (x-axis). Each genomic region has been labelled and categorized on the basis of the enrichment patterns. The percentages indicate the coverage of the region in the genome. (b) A schematic workflow of PAD for clustering DNA-binding proteins (e.g. transcription factors and chromatin remodellers) and histone modifications at functional genomic regions generated by ChromHMM.

Previous studies have largely focused on the role of LSD1 at enhancers^6, 27^, and despite the pervasiveness of LSD1 binding at promoters^27^, its role at promoters remains largely uninvestigated. To investigate the role of LSD1 at promoters, we correlated its binding profiles with other DNA- and chromatin-binding proteins at promoter regions using PAD. Our analysis shows a strong co-localization of LSD1 at promoters with core RNAPII pausing factors (RNAPII, CDK9, NELF-A, and SPT5) and additional factors such as MYC^3, 33, 34^ and BRD4^35–38^ (Fig. 2a). In particular, LSD1 co-localized most strongly with CDK9, the enzymatic subunit of the pause release factor P-TEFb^39–41^ across the three genomic regions (Fig. 2a). When we further investigated the co-localization at the binding sites of LSD1, we found that while the ChIP-seq signal of LSD1-CoREST complex components, CoREST, HDAC1, and HDAC2, were observed at comparable levels at promoters, bivalent domains and enhancers (Fig. 2b), the co-localization of LSD1 with RNAPII pausing and release factors was the strongest at promoters (Fig. 2b).

**Figure 2.**
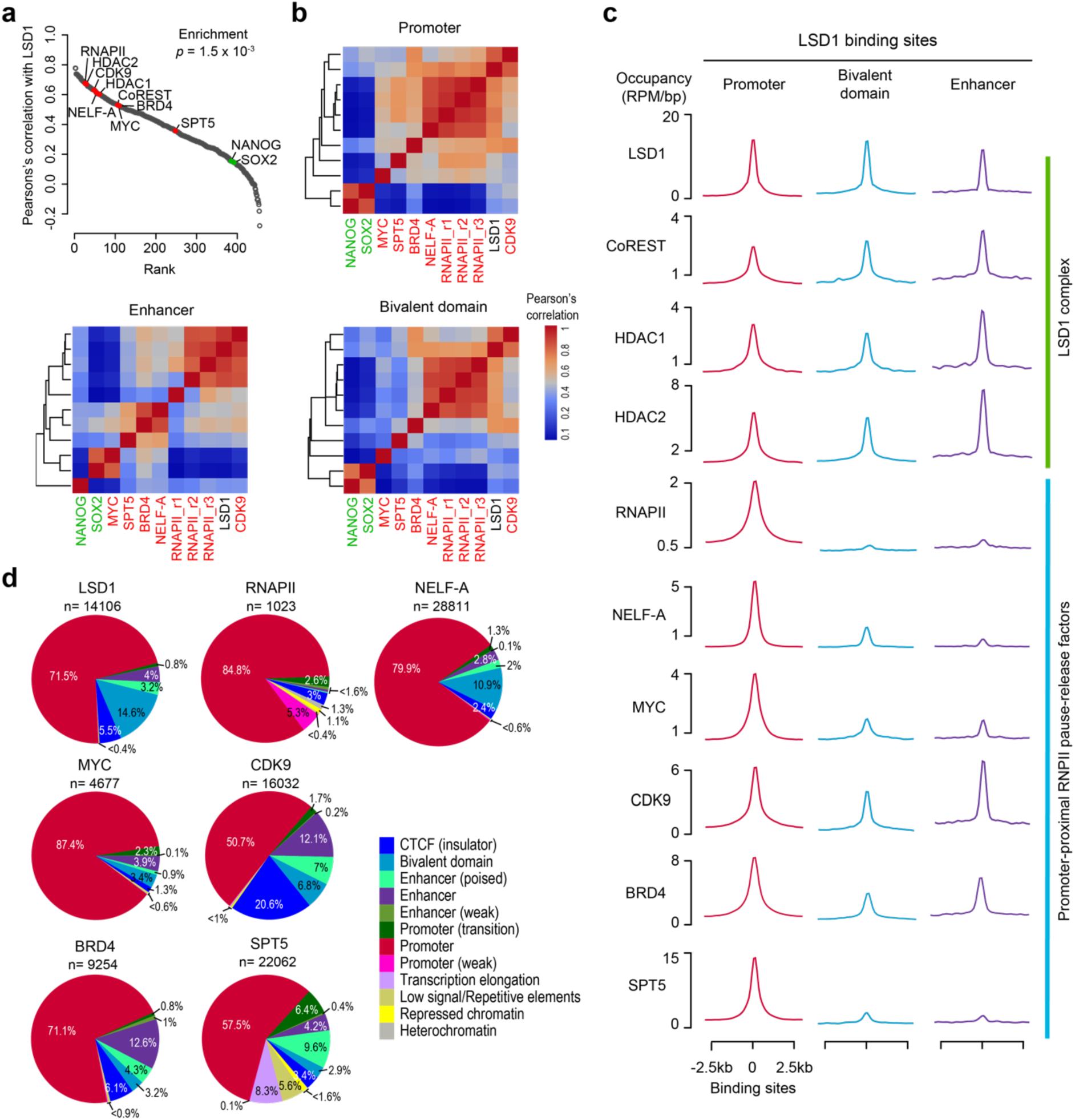
Co-localization of LSD1 with RNAPII pausing machinery across regulatory regions. (a) DNA binding proteins are ranked by the correlation of binding sites with LSD1 at promoter regions. A panel of genes known to be associated with RNAPII pausing are highlighted in red and two ESC pluripotency associated TFs are highlighted in green for contrast. Gene set enrichment test was applied to the RNAPII-associated proteins (those in red) with respect to their correlation with LSD1. (b) Co-localization heatmaps of LSD1 and select of RNAPII associated factors across three regulatory regions (promoters, bivalent domains, and enhancers). (c) Binding sites of LSD1, components of the LSD1 complex (CoREST, HDAC1, and HDAC2), and factors associated with the RNAPII pausing machinery (RNAPII, NELF-A, MYC, CDK9, BRD4, and SPT5) at the three regulatory regions. (d) Pie charts showing the proportion of binding sites of factors across the 12 regulatory regions.

We next assessed the binding profile of LSD1 (n=14106) across the genome and found that approximately 71.5% of all LSD1 binding sites in ESCs are found at the promoters (Fig. 2c). This was akin to the strong promoter-proximal binding preference of other RNAPII pausing and release factors (Fig. 2c). The prevalence of LSD1 binding at promoters is striking particularly given that previous studies have largely focused on the role of LSD1 at enhancers^27^, even though less than 8% of LSD1 binding sites fall into the enhancer regions (as defined by ChromHMM). Altogether, these data show strong recruitment of LSD1 to gene promoters and its co-localization with factors associated with RNAPII pausing.

### LSD1 Regulates the Transcription of Genes Associated with RNAPII Pausing

Given the widespread presence of LSD1 at gene promoters, we first investigated the relationship between gene expression and LSD1 binding. We found that the level of LSD1 binding at gene promoters correlated positively with gene expression in ESCs (Fig. 3a), which is consistent with experimental studies that have shown LSD1 is a transcriptional activator^42, 43^. We next sought to assess the relationship between RNAPII pausing and LSD1 binding of genes. To do this, we calculated the RNAPII pausing index (PI)^44^ of all genes based on RNAPII ChIP-seq data (see Methods) and correlated them with LSD1 ChIP-seq signal. Our analysis suggests that LSD1 is preferentially bound at the promoters of paused genes (Fig. 3b). Furthermore, we found that LSD1-bound genes show similar levels of RNAPII PI to those bound by other factors known to regulate RNAPII pausing or release in ESCs (Fig. 3c). These results suggest that LSD1-bound genes are subject to RNAPII pausing.

**Figure 3.**
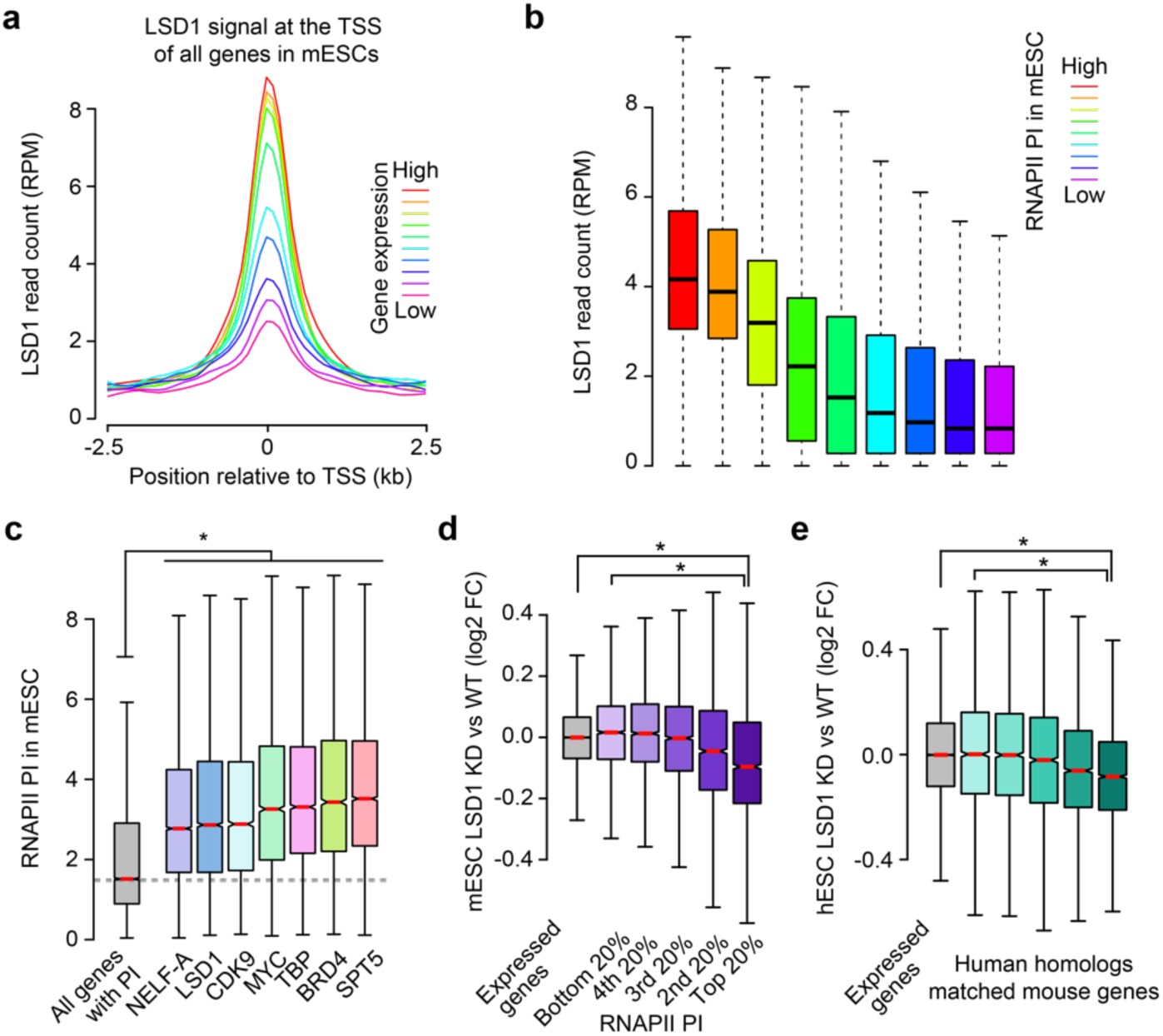
Knockdown of LSD1 affects genes with higher RNAPII pausing than those with lower pausing. (a) Gene sets were partitioned according to the level of expression in ESCs and the level of LSD1 signal (RPM) (+/− 1kb around the at the transcription start site [TSS]) were quantified for each gene set. (b) Boxplot of LSD1 signal (RPM) at gene promoters grouped according to RNAPII pausing index (PI), calculated as the ratio of RNAPII binding across the gene body against RNAPII binding at the TSS (see Methods). (c) Boxplot of RNAPII PI of genes bound by NELF-A, LSD1, CDK9, MYC, TBP, BRD4, and SPT5. (d, e) Boxplot of log2 fold-change in gene expression for gene sets grouped in terms of RNAPII PI after knockdown of LSD1 in (d) ESCs and (e) human ESCs (hESCs). * denotes statistical significance (p < 0.05) using Wilcox rank sum test.

We reasoned that if LSD1 has a functional role in controlling gene expression via regulation of RNAPII pausing, loss of LSD1 expression should preferentially affect the expression of LSD1 target gene sets with higher RNAPII PI much more so than those with lower RNAPII PI. To test this, we partitioned all quantified genes in ESCs into five groups and assessed their log2 fold change in expression before and after LSD1 knockdown^45^. We found that the log2 fold change in gene expression after LSD1 knockdown is proportionately affected by RNAPII pausing, indicating that LSD1 knockdown preferentially affects genes with higher RNAPII PI (Fig. 3d). Lastly, we confirmed our observations using data from human ESCs by showing the same pausing-dependent log2 fold change in expression with LSD1 knockdown can be observed in mouse and human ESCs^46^ (Fig. 3e). Collectively, these results suggest a potential role for LSD1 in regulating gene expression through RNAPII pausing, and these findings motivated us to investigate the mechanistic role for LSD1 in modulating transcriptional programs associated with RNAPII pausing.

### LSD1 and MYC Co-occupied Sites are Enriched for RNAPII Pausing Factors and Their Target Genes Show More Pausing

MYC is known to play a major role in RNAPII release in mammalian cells^3^. It binds to promoter-proximal regions of transcribed genes to regulate release of paused RNAPII by recruiting P-TEFb^3^. Recently, MYC has also been shown to recruit the elongation factor SPT5, a subunit of the elongation factor DSIF, to promoter-proximal RNAPII, wherein SPT5 travels with RNAPII and enhances its processivity during transcriptional elongation^34^. Given the important role of MYC in regulating transcriptional activity of RNAPII and the strong co-localization between LSD1 and the pause-release factor P-TEFb, we asked whether comparing sites of LSD1 and MYC co-localization against sites where either LSD1 or MYC are found alone may provide additional insight into the role of LSD1 in RNAPII pause release.

To test this, we partitioned bindings sites of LSD1 and MYC into three categories: MYC-specific sites, co-localized sites, and LSD1-specific sites (Fig. 4a). Approximately 57% of MYC binding sites were co-localized with LSD1 and 19% of LSD1 binding sites were co-localized with MYC. MYC-specific and LSD1-specific sites are significantly depleted of LSD1 and MYC, respectively, whilst the co-localized sites had comparatively similar amount of factor when compared to their respective individual sites (Fig. 4b).

**Figure 4.**
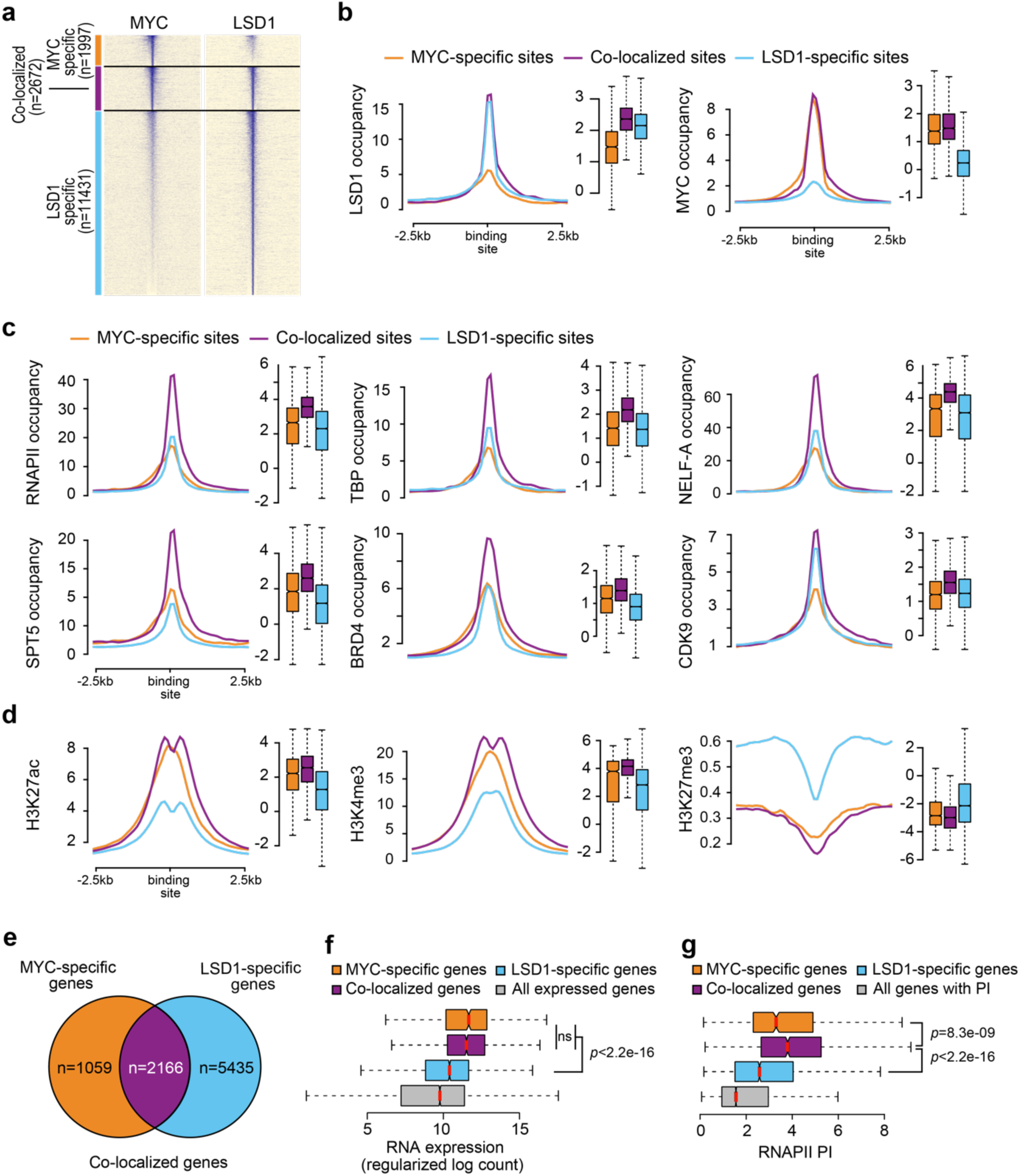
Signal density of factors at the binding sites of MYC-specific, co-localized, and LSD1-specific sites. (a) Heatmap showing the overlap of binding sites by the enrichment of either MYC or LSD1. (b) Density plots and boxplots of MYC and LSD1 occupancy (RPM) at the three genomic sites. The three lines denote the averaged signal across sites from each category: MYC-specific (orange), co-localized (purple), and LSD1-specific (blue) sites. Density plots and boxplots of (c) RNAPII machinery and pausing factor occupancy (RPM) and (d) histone modifications (RPM) at the three genomic sites. The ChIP-seq density plots were generated by calculating the number of reads within ± 2.5 kb upstream and downstream of the binding sites in 100 bp windows and normalized to RPKM. All boxplots show significant differences unless otherwise indicated. Wilcoxon rank-sum test was used in (b-d). (e) Venn diagram showing the overlap of binding sites by presence of either MYC and/or LSD1. Boxplots of (f) regularised and log-transformed RNA expression and (g) RNAPII pausing. Wilcox rank-sum test was used in (f, g).

Next, with the hypothesis that LSD1 co-occupancy would show us a differential binding profile of RNAPII pausing factors at MYC binding sites, we evaluated the difference in occupancy of RNAPII pausing factors (RNAPII, TBP, NELF-A, SPT5, BRD4, and CDK9) at these regions. Strikingly, we saw that compared to the LSD1/MYC-specific sites, sites co-occupied by LSD1 and MYC were significantly enriched by all RNAPII pausing factors. Interestingly, we observed that MYC-specific and co-localized sites are epigenetically more permissive with higher H3K27ac and H3K4me3 and lower H3K27me3 signals (Fig. 4d) than LSD1-specific sites, suggesting their target genes may be more transcriptionally active. Collectively, these findings show that MYC and LSD1 co-localized sites are active promoters that are enriched for RNAPII pause/release factors.

To further characterize genes regulated by either LSD1 or MYC or both (Fig. 4e), we identified the target genes bound by LSD1 and/or MYC and analyzed their expression profiles in ESCs. Whilst all three gene sets have higher than average transcription in ESCs (Fig. 4f), we found that genes targeted by MYC or both MYC and LSD1 were significantly more expressed than LSD1-specific target genes (Fig. 4f), consistent with what we observed at the epigenetic level (Fig. 4c, d). We next compared the RNAPII PI of the three sets of target genes (Fig. 5g). Consistent with our observation that the co-occupied sites are enriched for RNAPII pausing factors, we found that genes co-occupied by LSD1 and MYC exhibited significantly higher RNAPII pausing than those with LSD1 or MYC alone. Again, to our surprise, we found that genes bound specifically by MYC, whose role in RNAPII pause release has been well-documented^3, 33, 34, 38, 47^, had significantly lower RNAPII pausing than those co-bound, further suggesting that co-localization of LSD1 and MYC is associated with greater RNAPII pausing.

**Figure 5.**
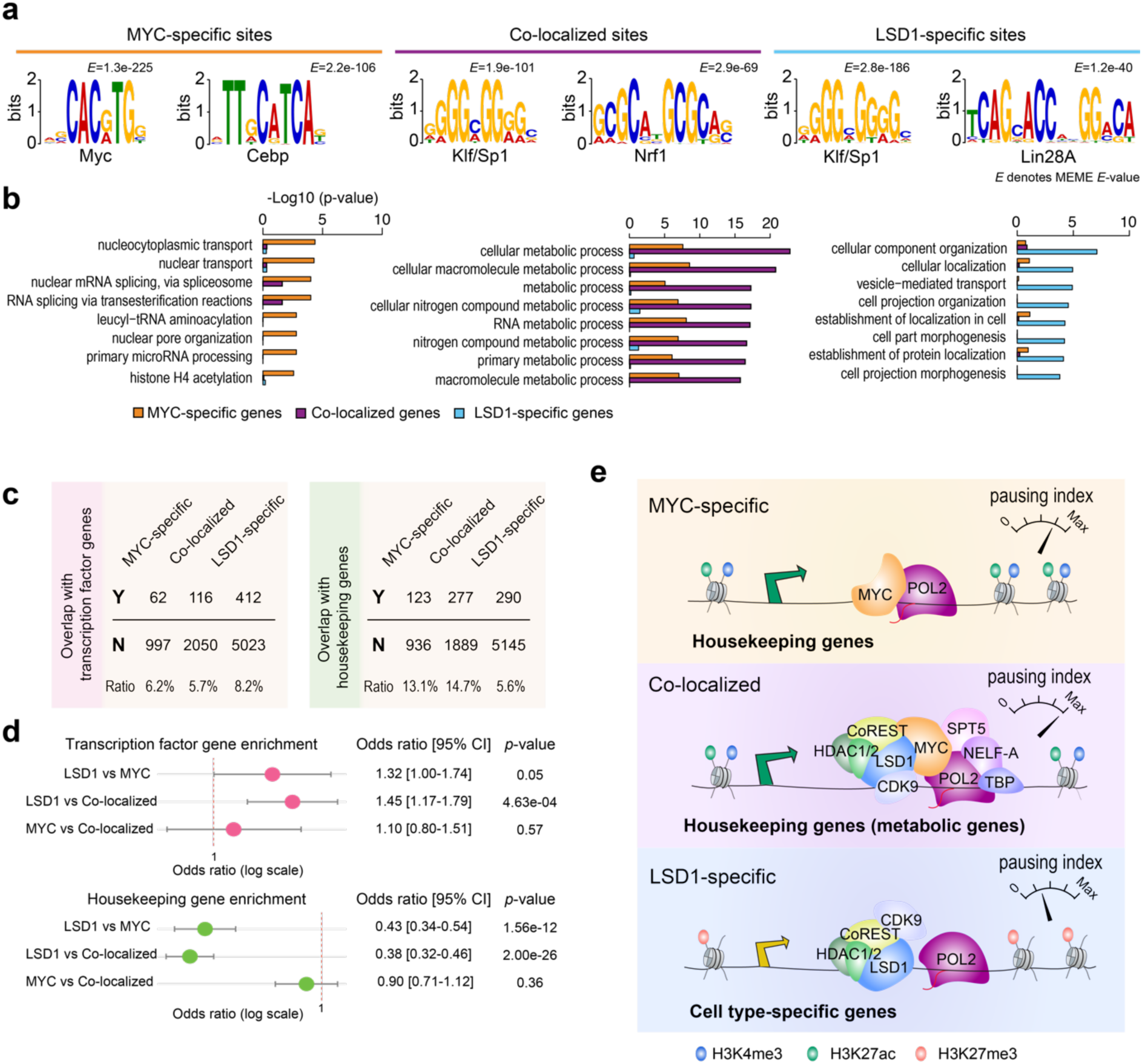
Differentially enriched of transcription factors and housekeeping genes at LSD1 sites. (a) Motif analysis with MEME for three categories of binding sites described in Fig. 4a. The y-axis denotes the sequence similarity independently of the query sequence length and database size, a higher score denotes a higher significant match. E-value or the expectation value refers to the expected hits of similar quality that could be found by chance, meaning the smaller the value the smaller the probability that the sequence can be found by chance. (b) Over-representation analysis of gene set enrichment from the three categories. X-axis denotes the degree of enrichment in terms of the negative log10 p-value. (c) Overlap and ratios of MYC, LSD1 and their co-localization sites with transcription factors (left) and housekeeping genes (right). (b) Test of non-independent overlap of (d) using Fisher’s exact test. (e) Schematic of the differential regulation of LSD1 target genes. Sites occupied by LSD1 only and those co-occupied with RNAPII pausing machinery show differential transcriptional control, epigenetic landscape and enrichment of transcription factors and housekeeping genes in ESCs.

### Occupancy of LSD1 and MYC is Associated with Different Gene Functions

Because our analysis suggested differential regulation of genes targeted by both LSD1 and MYC vs. those targeted by LSD1 or MYC alone, we performed motif discovery from the DNA sequences of the three sets of binding sites and gene set enrichment from the three gene sets to further characterize their difference (Fig. 5a, b). We found that the top two enriched motifs differed between the three sets of binding sites. Specifically, we found that the sites co-occupied by LSD1 and MYC, but not those by MYC alone, were enriched with motifs of KLF family member SP1^48^ (Fig. 5a). This is in agreement with previous research that found that KLF family members are DNA-binding transcription factors specifically associated with RNAPII promoters with slow TBP (TATA-binding protein) turnover, which is associated with high transcriptional activity, when compared to promoters with fast TBP turnover^49^. Lastly, we demonstrate that different gene sets are enriched in each category. Whereas MYC-specific genes are enriched for pathways such as RNA splicing and miRNA processing, in agreement with previous knowledge^50^, co-localized genes are largely enriched for metabolic processes, and LSD1-specific genes are enriched for pathways pertaining to cellular organization and localization (Fig. 5b). Together, these results suggest that genes co-bound by LSD1 and MYC and those bound by them individually may undergo different transcriptional regulation and perform distinct biological functions.

Given the different functional enrichment of genes found by the above analysis, we next conducted a test for non-independent overlap of the three gene sets with transcription factors, which are frequently associated with enhancers and cell type-specificity^51^, and housekeeping genes. Intriguingly, we found a reciprocal pattern of enrichment for transcription factors vs. housekeeping genes in the three gene sets. LSD1-specific target genes were enriched for target genes encoding transcription factors whilst MYC-specific and co-localized genes were not (Fig. 6a, b). In contrast, co-localized and MYC-specific target genes were enriched for target genes encoding housekeeping genes whilst LSD1-specific genes were depleted, consistent with the gene set enrichment results showing that these sites were enriched for genes related to, for example, metabolic processes that take place regardless of the cell type (Fig. 5e). Collectively, our findings implicate a putative role in pausing of LSD1 at promoters through its association with MYC and P-TEFb that may be distinct from its enhancer-related role in ESC differentiation^27^ and demonstrates differential transcriptional control, epigenetic landscape, and enrichment of transcription factors and housekeeping genes between LSD1 sites distinguished by RNAPII pausing (Fig. 6c).

## Discussion

Here we show how the association of LSD1 at the promoters of actively transcribed genes corresponds to genes engaged in RNAPII pausing in ESCs and implicates a functional role of LSD1 in RNAPII pausing. LSD1 has been shown to decommission enhancers during ESC differentiation^6, 27^, but its role at the promoters of genes remains elusive. Several lines of evidence suggest the involvement of LSD1 in RNAPII pausing. We observed that LSD1 signal was strongly and positively correlated with gene expression, supporting a transactivating role of LSD1 in gene transcription. This is consistent with a recent study that demonstrates that LSD1 is recruited to promoters by MYCB to promote active transcription^52^. Previously, LSD2 the only mammalian homolog of LSD1 has also been shown to have an important function in active gene transcription elongation^53^. Moreover, our genome-wide chromatin-binding profiles demonstrated that LSD1 binding sites at RNAPII paused sites are strongly co-localized with CDK9 (Fig. 2b, c), the enzymatic subunit of P-TEFb, whose activity has been proposed as the rate-limiting step in paused RNAPII release^39, 54, 55^. The co-localization of LSD1 and CDK9 was observed at both promoter and enhancer elements, where CDK9 have been shown to be involved in enhancer-associated RNAPII pausing^17, 56, 57^. A recent study has demonstrated a functional role of SIRT6, a histone deacetylase, in the release of RNAPII pausing by preventing the eviction of NELF-E from the chromatin^58^, highlighting the direct involvement of chromatin remodellers in RNAPII pausing and release.

Previous studies have shown that LSD1 physically interacts with RNAPII^59^, MYC^34, 60^, and SNAIL^61^, which in Drosophila embryos has been shown to inhibit the release of paused RNAPII^62^. Among 22 methyltransferases, LSD1 was the only one found to be in direct physical contact with MYC in the methyltransferase interactome^34, 60^. A recent study demonstrated a generic role of MYC in governing the transition of RNAPII from a paused state into a transcriptionally engaged mode by directly recruiting SPT5 to RNAPII^34^. Consistent with the function of MYC as a universal amplifier of gene expression^63^, the ablation of MYC led to a global depletion of SPT5 in chromatin^34^. Strikingly, we found that among the MYC binding sites, the presence of LSD1 facilitated the enrichment of RNAPII pausing factors at the promoters of genes and a higher pausing index (Fig. 4). In comparison, genes bound only by MYC demonstrated lower RNAPII pausing index but still retained a transcriptional level comparable to genes bound by both factors. Consistent with these findings, it has previously been shown that genetic ablation and chemical inhibition of LSD1 in pluripotent cell lines (mouse ESCs and F9 cells) increased the level of H3K56ac^64^, a histone modification associated with the gene body^2^ and recently identified as a key histone mark, along with H3K9ac, that modulates transcriptional pausing and elongation^58^. In another study, LSD1 ablation and catalytic inhibition led to an upregulation of H4K16ac^64^, a histone mark that has been implicated in the regulation of RNAPII promoter-proximal pausing by recruiting BRD4 and P-TEFb^65, 66^. Thus, our integrative analysis suggests a role for LSD1 in the regulation of RNAPII pause release through its association with critical pause release factors MYC and pTEFb; however, functional and mechanistic studies remain to be carried out to validate how LSD1 regulates RNAPII promoter-proximal pausing.

In ESCs, the catalytic activity of LSD1 has been shown to render enhancers inactive and is associated with the silencing of target genes^27^. Studies have shown LSD1 occupies the regulatory domains of pluripotency genes, including transcription factors, and decommissions them only when cells undergo lineage-specific differentiation^6, 27^. While we found that LSD1-specific genes are enriched for transcription factors, supporting its role in regulating cell type-specific differentiation, genes co-localized by LSD1 and MYC are significantly depleted of transcription factors and rather enriched for housekeeping genes, and in particular those that are involved in the regulation of cell type-invariant metabolic processes. Consistent with this, loss of LSD1 function has been shown to be associated with significant proliferative defect in neural^67^ and embryonic stem cells^64^ and has been implicated in cellular growth pathways and to the metastatic and oncogenic potential of several types of cancer^28, 42, 68^. Collectively, our integrative analysis presents evidence for a previously unanticipated role of the histone demethylase LSD1 in cooperation with other RNAPII pausing factors in modulating cell type-specific and cell type-invariant genes.

## Methods

### ChIP-seq data analysis

All ChIP-seq data analyzed in this study were generated from mouse ESC lines. Reads from each ChIP-seq dataset were aligned to the mouse genome (mm9 assembly) using Bowtie version 0.12.8^69^. Reads that mapped to unique genomic regions with no more than two mismatches were retained for analysis. For each DNA-binding protein, mapped reads were binned across the mouse genome with 1kb bin size and quantified as reads per base/kilobase per million reads (RPKM) and used in PAD clustering (see next section). ChIP-seq read density plots were generated by calculating the number of reads within ± 2.5 kb upstream and downstream of sites of interest in 100 bp windows and normalized to RPKM and plotted as histograms. Data for heatmaps were generated in a similar manner.

### PAD implementation

PAD2 was developed in Python 3.7 based on Django web framework and the interactive plots in PAD2 were rendered using Plotly.js, an extension from PAD clustering^31^ implementation. PAD clustering^31^ (http://pad2.maths.usyd.edu.au) was previously developed to characterise co-localization of TFs and epigenomic marks at various genomic regions based on their ChIP-seq profiling. Here, we extended PAD by increasing the number of partitions for genomic binding sites and histone modifications to 12 functional genomic regions identified by ChromHMM^32^ in ESCs (Fig. 1).

### Mapping of genome to functional regions

A collection of transcription factors, chromatin remodellers and histone mark ChIP-seq datasets have been processed and quantified in 1kb bin across the genome. For each of the peak files, we mapped them to the 12 genomic functional regions (Fig. 1) by using the intersect method in BEDTools v2.28.0^70^ and calculate their fold change.

The fold change of a protein binding at each functional region is defined as follows:

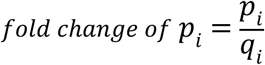

where *p_i_* denote the percentage of protein binding at genomics region *i*, and *q_i_* denote the percentage of genomic region *i* covers in a whole genome.

### Clustering of DNA binding proteins

To perform clustering of DNA binding proteins at different genomic regions, we combine a set of peak files from a user-specified selection of DNA binding proteins and functional regions to a matrix. We then perform hierarchical clustering with Pearson’s correlation as a similarity metric.

### Binding sites and target genes

To define binding sites for each DNA-binding factor, aligned reads were processed using SISSRs with a common input^71^ and a those with a *P* < 0.001, a stringent cut-off, were called as binding sites^72^. MYC-specific and LSD1-specific sites were defined as any MYC or LSD1 peaks that lack LSD1 or MYC ChIP-seq signals, respectively, at the same locus. Co-localized sites were determined as loci where MYC and LSD1 peaks overlap (within 500 bp window). Based on the mm9 RefSeq annotation, genes with closest TSSs to binding sites (within 1.5 kb) of a DNA-binding factor are assigned as the target genes of that factor.

### Calculation of RNAPII pausing index

To calculate the RNAPII pausing index, we used the method described previously^73^. Briefly, for each gene, we calculated the pausing index as follows:

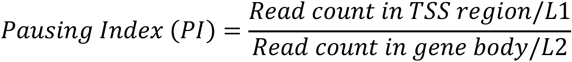

where the transcription start site (TSS) region of a gene is defined as the −50 bp to +300 bp around the TSS and the gene body is defined as +300 bp downstream of the TSS to +3 kb past the transcription end site. To segregate genes into different tiers of pausing, we categorized genes into five pausing groups on the basis of their RNAPII pausing index at the promoter so that each group contained roughly the same number of genes. The same pausing groups (i.e., homologous genes) were used to evaluate the effect of LSD1 knockdown in hESCs.

### Gene expression analysis

Previously published data from LSD1 knockout and WT mouse ESC samples measured using Illumina MouseWG-6 v2.0 expression beadchip were downloaded from the NCBI Gene Expression Omnibus (GEO; http://www.ncbi.nlm.nih.gov/geo/) with accession numbers GSE21131^45^. Log2 fold change for each gene was then calculated by averaging across duplicates in either LSD1 KD or WT samples and subtracting log2 transformed values in LSD1 KD measurement with WT measurement. The same approach was taken to process data from MYC knockdown and WT mouse ESC samples measured using Illumina HiSeq 2500 (GSE113329^74^) and microarray data from LSD1 knockdown and WT human ESC samples measured using Affymetrix Human Promoter 1.0R Array from human ESCs (GSE24844)^46^. RNA-seq data from ESCs in naïve state (GSE117896) was used for defining expressed genes^75^. Specifically, expressed genes were defined as any gene with regularised log expression equal or higher than 5.

### Motif enrichment analysis

For motif enrichment analysis, DNA sequences (500 bp) flanking the center of the binding sites for each factor were first extracted from mm9 assembly using the ‘getfasta’ of the bedtools^76^. These sequences were subsequent searched using MEME^77^ using a minimum and maximum window size of 5 and 15, respectively, and the zero or one motif occurrence per sequence (zoops) option. The top two most enrichment motifs (based on the MEME reported E-value) were presented and annotated using known motifs.

### Functional enrichment analysis

Functional enrichment analysis, in terms of overrepresentation of genes from a pathway, was performed using Fisher’s exact test against the gene ontology (GO) terms from the GO database^78^. P-values were adjusted using the Benjamini-Hochberg method.

### Statistics and reproducibility

Visualizations and statistical tests were performed in the environment of the R Project for Statistical Computing (https://www.r-project.org). For comparing statistics between two groups, Wilcox rank sum test was employed. P values were specific on each plot to show statistical significance.

## Data availability

The authors declare that all data supporting the findings of this study are available within the article and its Supplementary Information files or from the corresponding author upon reasonable request. The accession codes for previously reported ChIP-seq datasets are as follows: MYC (GSE11431); NANOG (GSE44286); Sox2 (GSE44286); SPT5 (GSE20530); NELF-A (GSE20530); RNAPII (GSE20530, GSE21917); BRD4 (GSE111264); LSD1 (GSE27841); CDK9 (GSE44286); TBP (GSE22303); CoREST (GSE27841); HDAC1-2 (GSE27841); H3K27ac (GSE117896); H3K4me3 (GSE117896); and H3K27me3 (GSE117896). The accession codes for previously reported microarray or RNA-seq data are GSE21131, GSE113329, GSE24844, and GSE117896.

## Authors’ contribution

P.Y. and H.J.K. conceived the study; H.J.K. and P.Y. performed bioinformatics analyses with contribution from T.K.; H.J.K. and P.Y. analyzed and interpreted the data with input from A.O.; H.J.K. and P.Y. wrote the manuscript. All authors reviewed, edited, and approved the final version of the manuscript.

## Acknowledgements

This work was supported by an Australian Research Council (ARC) Discovery Early Career Researcher Award (DE170100759) and a National Health and Medical Research Council Investigator Grant (1173469) to P.Y., an ARC Postgraduate Research Scholarship and Children’s Medical Research Institute Scholarship to H.J.K., the Judith and David Coffey Life Lab Scholarship to T.K. We thank our colleagues from the Embryology Unit, Children’s Medical Research Institute, for their discussion and feedback.

